# TaxaNorm: a novel taxa-specific normalization approach for microbiome data

**DOI:** 10.1101/2023.10.31.563648

**Authors:** Ziyue Wang, Dillon Lloyd, Shanshan Zhao, Alison Motsinger-Reif

## Abstract

**Background:** In high-throughput sequencing studies, sequencing depth, which quantifies the total number of reads, varies across samples. Unequal sequencing depth can obscure true biological signals of interest and prevent direct comparisons between samples. To remove variability due to differential sequencing depth, taxa counts are usually normalized before downstream analysis. However, most existing normalization methods scale counts using size factors that are sample specific but not taxa specific, which can result in over- or under-correction for some taxa.

**Results:** We developed TaxaNorm, a novel normalization method based on a zero-inflated negative binomial model. This method assumes the effects of sequencing depth on mean and dispersion vary across taxa. Incorporating the zero-inflation part can better capture the nature of microbiome data. We also propose two corresponding diagnosis tests on the varying sequencing depth effect for validation. We find that TaxaNorm achieves comparable performance to existing methods in most simulation scenarios in downstream analysis and reaches a higher power for some cases. Specifically, it has a well balance on power and false discoveries control. When applying the method in a real dataset, TaxaNorm has improved performance when correcting technical bias.

**Conclusion:** TaxaNorm considers correcting both sample- and taxon-specific bias by introducing an appropriate regression framework in the microbiome data, which aids in data interpretation and visualization. The ‘TaxaNorm’ R package is freely available through the CRAN repository https://CRAN.R-project.org/package=TaxaNorm and the source code can be downloaded at https://github.com/wangziyue57/TaxaNorm.

## Background

There is growing evidence that microbial communities influence human health [1, 2]. Advanced high-throughput sequencing technologies (HTS) such as 16S ribosomal RNA (rRNA) gene amplicon sequencing (16S) and whole-genome shotgun sequencing (WGS) allow researchers to survey microbial communities in a study population [3–7]. While HTS offers advantages in precision and accuracy, its use can be limited by sequencing depth (library size), which is the total number of reads obtained per sample from equipment [8–10]. The raw data is compositional and represents only a fraction of the species abundance in each sample from an ecosystem with unknown microbial volume and thus there can be significant variation in sequencing depth between samples, even within the same biological community [8, 11, 12]. Thus, the observed differential abundance (DA) between samples, which, in theory, reflects biological variation, is confounded by sequencing depth [13–15]. Therefore, data are usually normalized to eliminate bias introduced by sequencing depth and to reflect true biological heterogeneity [12, 16]. In sufficiently normalized data, taxa abundance should be independent of sequencing depth across samples.

Normalization approaches currently used for microbiome data can be broadly classified into three types - rarefaction, log-ratio transformation, and scaling. Rarefaction is commonly used in early-stage microbiome studies [17–19]. Reads are randomly drawn without replacement in each sample, such that all samples have the same total count and thus the same sequence depth [17]. A major limitation of rarefaction is the use of an arbitrary cut-off value across samples, resulting in a loss of statistical power and sample heterogeneity due to decreased sample size [12]. Log-ratio transformation is used to normalize compositional data by taking the log-ratios of all taxa with respect to a fixed reference component [20–24]. To handle the zeros commonly seen in microbiome data [15], an arbitrary pseudo count is typically used to replace zeros, but the choice of this arbitrary value can influence downstream analysis [25–27]. Further, the statistical inference is based on relative change with respect to the chosen reference. In order to recover the true scale of taxa and compare differences in the absolute counts, we focus on scaling in this manuscript.

Scaling is a common normalization approach that divides raw counts by a sample-specific size factor across all taxa. Algorithms to estimate size factors include total-sum scaling (TSS), which simply scales samples by their sequencing depth, median-by-ratio (MED) from DESeq2 [28], upper quartile (UQ) [29] and trimmed mean of M-values (TMM) [30] from edgeR [31], cumulative sum scaling (CSS) from metagenomeSeq [15], Wrench [32], and analysis of compositions of microbiomes with bias correction (ANCOM-BC) [11]. A major drawback of most scaling methods is the use of a common size factor to represent sequencing efficiency, which is the effect of sequencing depth on all taxa in a given sample. In practice, however, there is evidence that sequencing efficiency varies across taxa, as a particular taxon may be preferentially measured during sequencing due to polymerase chain reaction amplification efficiency or other technical reasons [15, 33–36]. For example, sequencing efficiency varies for gram-negative and gram-positive bacteria because it is more difficult to extract DNA from gram-positive bacteria during sample collection due to their strong cell walls. As a result, gram-positive bacteria may be under-represented in observed taxa abundance [37].

Figure 1 provides an example from a human gut microbiome dataset of 510 taxa from 300 healthy individuals. The dataset is described in detail in the real-data application section. We examined the relationship between the counts and sequencing depth for each taxon. Two specific taxa, *Dehalobacterium* and *Bacteroides*, were shown as an example. The monotonic increasing trend in the raw counts differed (Figure 1a), and neither TSS nor ANCOM-BC had good performance as they led to over- or under-correction of the effect of sequencing depth (Figure 1b, 1c). We further regressed all taxa counts on sequencing depth individually from the above data using zero-inflated negative binomial (ZINB) regression and compared the corresponding coefficients for each taxon. Under the assumption of scaling methods, coefficients should be fixed and nearly identical. To better visualize the results, we combined the taxon-specific coefficients by their phylum group. However, density plots indicate that coefficients differ by taxa (Figure 2), suggesting that the relationship between taxa count and sequencing depth varies across taxa. The coefficient distribution was not specific to the example we showed and was generalized to other microbiome datasets (Supplementary Figure 7, 8) [38, 39].

**Fig. 1.**
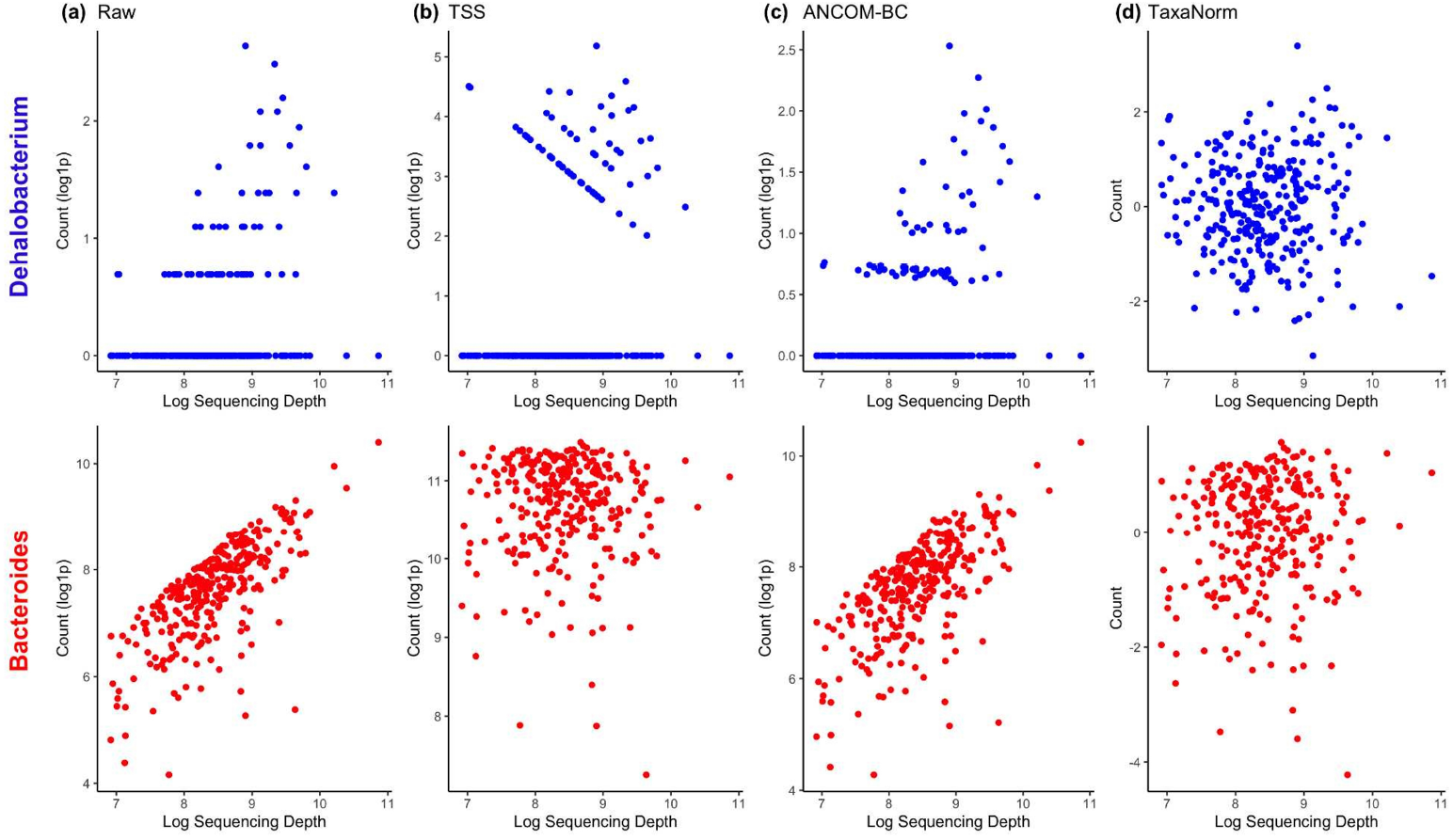
Relationship between counts and sequencing depth before and after normalization. Two microbial organisms, *Dehalobacterium* (blue) and *Bacteroides* (red), are presented as examples. (a) Raw counts. (b) Normalized counts by TSS. (c) Normalized counts by ANCOM-BC. (d) Normalized counts by TaxaNorm. Each stool sample is represented by a single dot. The sequencing depth (number of reads) is shown in log-scale. The raw count in panel (a) and normalized count via TSS (b) and ANCOMBC (c) are shown with log1p transformation with a pseudo number of 1 to avoid undefined values for log(0). The normalized counts from TaxaNorm (d) are shown in raw scale.

**Fig. 2.**
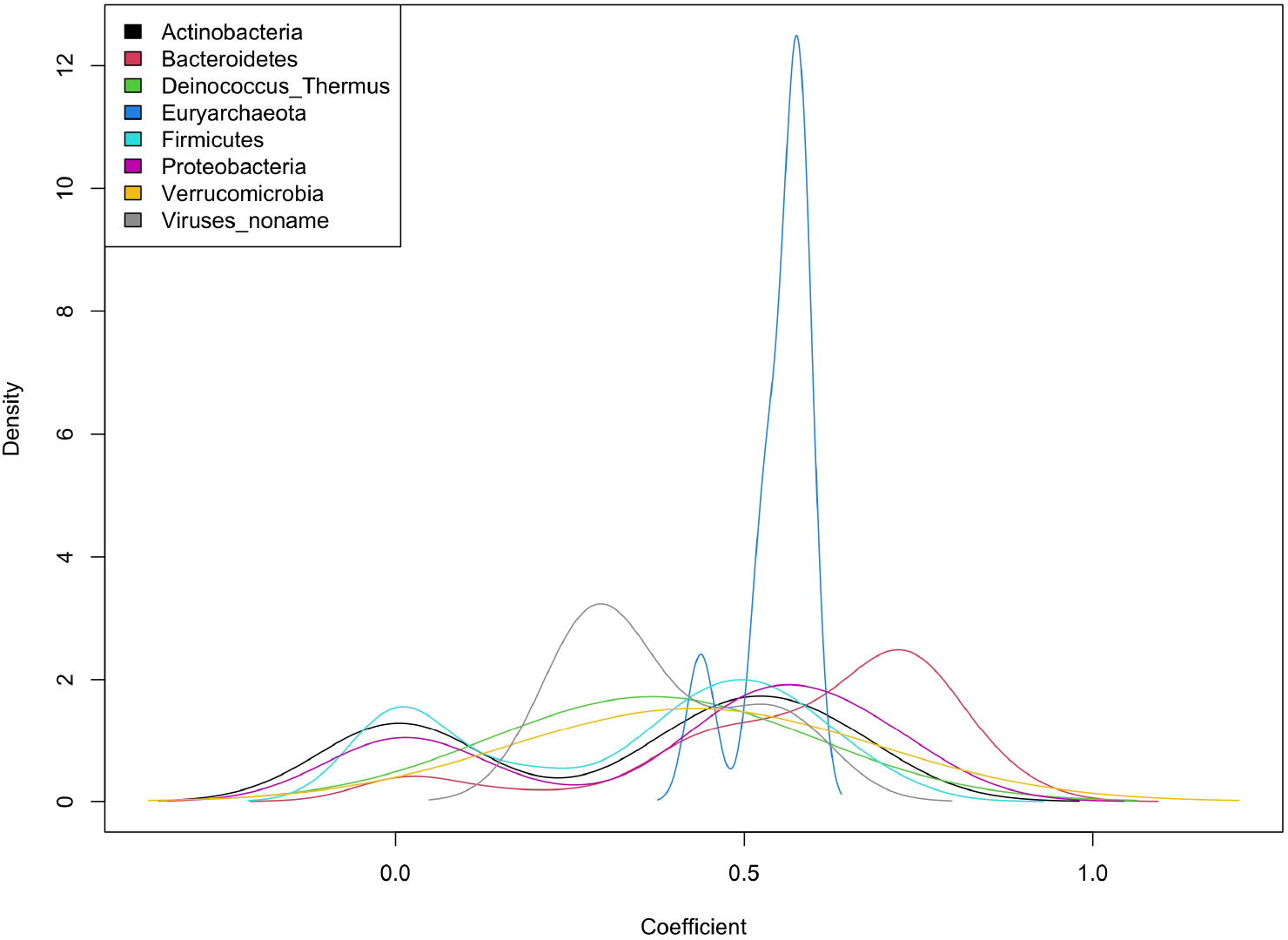
Sequencing efficiency of each taxon in human gut microbiota data. The relationship between raw taxa-specific counts and sequencing depth was estimated using ZINB regression. Densities of corresponding coefficients are colored by phylum rank for all taxa.

Additionally, high heterogeneity has been observed within samples in microbiome data [40, 41]. Although CSS and Wrench consider a taxon-specific factor, they rely on zero-inflated Gaussian distribution where the sample heterogeneity will be missed. Consequently, the choice of normalization method should be not only sample-specific but also taxon-specific and based on a model that adequately captures the nature of microbiome data.

Motivated by these observations, we developed TaxaNorm, a novel normalization method based on a ZINB model, that allows the effects of sequencing depth on taxa abundance to vary by microbial organism. TaxaNorm can handle both structural (biological) and sampling zeros. Further, this method allows the magnitude of over-dispersion to depend on sequencing depth for microbiome data. In contrast to traditional fixed-dispersion negative binomial models such as DESeq2 and edgeR, TaxaNorm includes a sequencing depth-dependent dispersion parameter to account for sample heterogeneity. The output from TaxaNorm can be used for variable selection, dimensional reduction, clustering, visualization, and differential abundance analysis.

## Materials and methods

The overall workflow of TaxaNorm is depicted in Supplementary Figure 1. Each step of the algorithm is detailed below.

### Varying-dispersion zero-inflated negative binomial model

We assume the observed taxa counts follow a ZINB distribution, which is a mixture of a negative binomial (NB) distribution of counts and a mass distribution at zero. The excess of zeros in microbiome data are handled in two ways: structural (biological) zeros through the mass distribution at zero and sampling zeros through the NB distribution. For a given taxon *i* (*i* = 1, …, *p*) in sample *j* (*j* = 1, …, *n*), let *Y*_*ij*_ ∼ **ZINB**(*μ*_*ij*_, *θ*_*ij*_, *π*_*ij*_) denote the observed counts so that

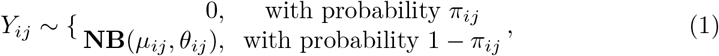

where *μ*_*ij*_ and *θ*_*ij*_ are the mean and dispersion of the NB distribution and *π*_*ij*_ is the probability of zero mass, or the called zero-inflation parameter. Under this parameterization, the variance of NB distribution is 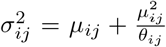. In particular, this NB distribution converges to a Poisson distribution when *θ*_*ij*_ → ∞.

To account for both sample- and taxon-specific effects of sequencing depth on counts, for a given taxon *i*, we have

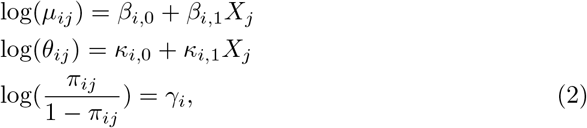

where *X*_*j*_ = log(Σ_*i*_ *y*_*ij*_) is the log of sequencing depth for sample *j* = 1, …, *n*. This formulation allows for the taxon-specific impact of sequencing depth on mean count (*β*_*i*,1_) and dispersion (*κ*_*i*,1_) to better capture the between-taxa variation and high heterogeneity in microbiome data compared to existing methods. Although many experimental and biological factors are linked with true zeros, we assume the zero-inflation parameter *π*_*ij*_ is taxon-specific only and is common across samples (*j*) since numerical evidence shows simpler models tend to have better model-fitting performance [42, 43]. Under the special case that *β*_*i*,1_ (*i* = 1, …, *p*) is equal to 1 for all taxa and *κ*_*i*,1_ = 0 (*i* = 1, …, *p*), TaxaNorm operates under a similar model assumption as most scaling normalization methods. On the other hand, when *κ*_*i*,1_ = 0 (*i* = 1, …, *p*), this is similar to the concept behind CSS and Wrench that dispersion does not change with sequencing depth.

### Parameter estimation

We fit the model and estimate the parameters for each taxon individually. For taxon *i*, given the observed counts *Y*_*i*_ = {*y*_*ij*_, *j* = 1, …, *n*}, and the sequencing depth *X* = {*x*_*j*_, *j* = 1, …, *n*}, the log-likelihood can be written as follows:

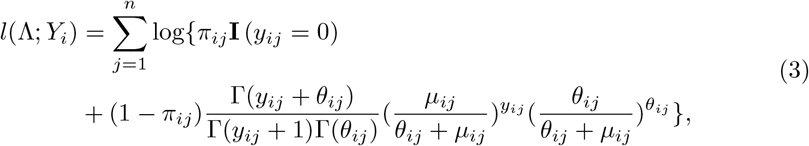

where Λ = (*β*_*i*,0_, *β*_*i*,1_, *γ*_*i*_, *κ*_*i*,0_, *κ*_*i*,1_) denotes the full set of unknown parameters in (2), and **I** (*·*) is an indicator function.

In practice, directly maximizing (3) causes difficulty when distinguishing zeros from the NB part and the zero-inflation part, leading to an unreasonably low estimation of *π*_*ij*_ [42]. Therefore, we used the expectation-maximization (EM) algorithm to obtain the maximum likelihood estimations (MLEs) 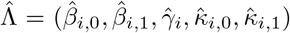. We defined a latent random variable, *Z*_*ij*_, to indicate whether *Y*_*ij*_ is generated from the zero mass (*Z*_*ij*_ = 1) or NB count (*Z*_*ij*_ = 0). The log-likelihood now becomes

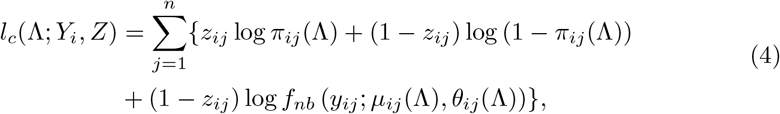

where *f*_*nb*_ denotes the probability mass function (PMF) of the NB distribution.

We set the starting values 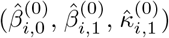 at the estimates from a ZINB regression with *κ*_*i*,1_ = 0, using the R built-in function *zeroinfl()* from the *pscl* package. For 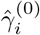, we initialized from a logistic regression with *Y*_*i*_ = 0 as the outcome to avoid the local maximum at the starting point [42].

### E step

For the *t* th iteration, the conditional expectation of the log-likelihood given the observed data *Y*_*i*_ and current parameter estimate 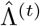 is computed as:

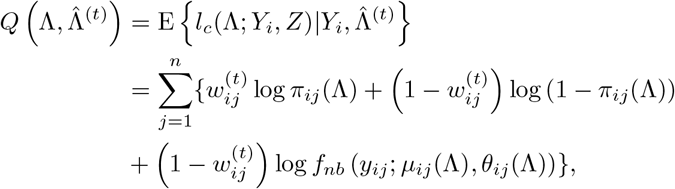

where

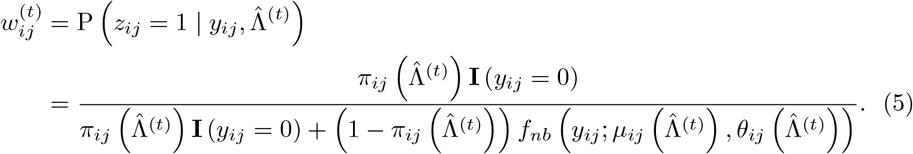

### M Step

The parameter estimate is updated as 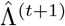, maximizing the quantity 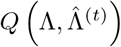:

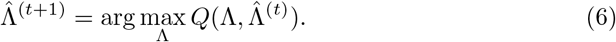

These two steps are repeated until convergence is achieved:

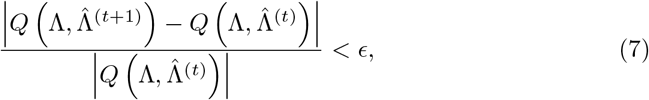

where *ϵ* is a small value threshold (here, 1*e* − 5).

Finally, the estimated 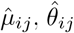, and 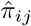 are calculated by plugging in 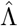 to the above regression model (2).

### Taxa-specific normalization and diagnosis testing

Using the MLEs 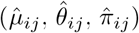, we calculated the quantile residuals [44] as normalized taxa counts to remove the effects of sequencing depth:

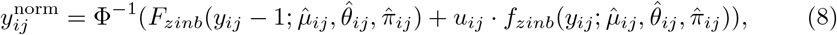

where Φ is the cumulative distribution function (CDF) of the standard normal distribution, *F*_*zinb*_ and *f*_*zinb*_ denote the CDF and PMF of the ZINB distribution, and *u*_*ij*_ is a random variable from a uniform distribution on (0, 1). Positive residuals for a given taxon in a given sample indicate greater abundance than expected given the taxon’s average abundance in the microbial community and sequencing depth while negative residuals indicate less abundance.

Correctly specifying the model is critical to increase power for any analysis. Therefore, we propose two model diagnostic tests. First, we test the existence of a sequencing depth effect on taxa counts from two perspectives - mean and dispersion (the ‘prevalence’ test):

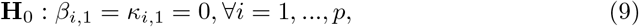

where the alternative hypothesis (**H**_*A*_) indicates that a sequencing depth effect exists for at least one taxon through one parameter. To do so, we use the likelihood ratio test (LRT). Specifically, under the null hypothesis, we fit a reduced intercept-only ZINB regression with fixed dispersion, where sequencing depth does not influence the taxa abundance. For this global test for all taxa, the likelihood ratio statistic is asymptotically, *χ*^2^, distributed with a degree of freedom of 2*p*.

Additionally, two major improvements of TaxaNorm are that it assumes the effect of sequencing depth is taxon-specific (i.e., differential sequencing efficiency), and the dispersion parameter depends on sequencing depth, while most scaling methods basically assume *β*_1,1_ = … = 1, *κ*_1,1_ = … = 0. To test whether TaxaNorm better reflects the data than existing scaling methods, we conduct an ‘equivalence’ test with the following null hypothesis

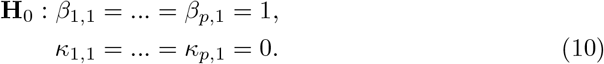

Again, we use LRT, where the MLE under the null hypothesis is estimated by restricting the equal effect of sequencing depth to be 1 on taxa abundance via the mean parameter, and fit a fixed dispersion ZINB regression. In this case, the likelihood ratio statistic is also asymptotically, *χ*^2^, distributed with a degree of freedom of 2*p*.

### Simulation studies

For all simulations, we set the true values of regression parameters as those estimated from a subset of 147 stool samples in a real microbiota dataset from the human microbiome study detailed in the real-data application section, which ensured our simulated data have similar characteristics as actual case-study data. The dataset comprises 510 taxa with non-zero counts observed in more than 10 samples.

For a given sample size *n*, we first randomly generated sequencing depth, *X*_*j*_ ∼ **U**(*a, b*), *j* = 1, 2, …, *n*, where *a* and *b* are the minimum and maximum sequencing depths in the template data. Together with the coefficients estimated from the template data 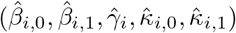, we calculated the mean 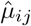, dispersion 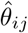, and zero mass 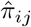 for each taxon across all samples based on model (2). We then generated taxa counts from ZINB distribution with these mean, dispersion, and zero mass parameters.

### Parameter estimation and diagnosis tests

We first assessed the performance of the proposed model diagnosis tests by considering scenarios with various coefficient settings. Specifically, we performed simulations under the two null hypothesis: 1) *β*_*i*,1_ = *κ*_*i*,1_ = 0 for all taxa, where sequencing depth effect does not exist for any taxa; and 2) *β* = 1, *κ* = 0, where the sequencing depth effect is the same across all taxa with fixed dispersion, to assess the type I error control for the tests proposed in (9) and (10). We conducted power analysis for prevalence tests by simulating various alternatives, where *β*_*i*,1_ = *κ*_*i*,1_ = 0 for a subset of taxa, while other parameters came from real data, so the sequencing depth effect exists for several taxa only. We also conducted power analysis for equivalence tests by restricting *β* = 1 for a subset of taxa, while other parameter values varied. We set the sample size from 100 to 1000 for all scenarios, and repeated each simulation scenario procedure 1000 times. See Supplementary Table 1 for details of the full simulation settings.

### DA taxa comparison

The efficiency of data normalization can be evaluated by assessing the influence of a method on downstream analysis. We conducted additional simulations under various settings to examine the performance of our proposed normalization method in detecting DA taxa using post-normalization counts.

To introduce DA taxa, we randomly assigned half of the samples to each group and manipulated the mean taxa count by multiplying the effect size. Specifically, we used a dummy variable, 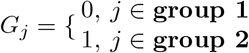, such that log(*μ*_*i,G*_) = *β*_*i*,0_ +*β*_*i*,1_*X*_*j*_ +*β*_*i*,2_*G*_*j*_. Here, *β*_*i*,2_ can be regarded as the log fold-changes (i.e.,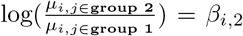). For example, we first allowed 50% of taxa to be DA, by randomly selecting 25% of all taxa with increased abundance in group 2 with the fold-changes (exp(*β*_*i*,2_)) generated from **U**(4, 10), and another 25% of randomly selected taxa with decreased abundance in group 2 (i.e.,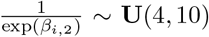). The remaining 50% of taxa were not DA (i.e., *β*_*i*,2_ = 0). We also investigated scenarios with varying percentages of DA taxa, namely 10% (5% increase and 5% decrease) and 20% (10% increase and 10% decrease), as well as smaller fold changes generated from **U**(2, 4). We varied sample sizes from 100 to 1000. The detailed simulation scenarios are outlined in Supplementary Table 2.

Next, we simulated a scenario without DA taxa, where the observed difference in counts was completely due to sequencing depth. For this scenario, we generated the sequencing depth of the two groups separately at different levels. The sequencing depth for samples in the second group was, on average, three times greater than for the first group (*X*_*j*_ ∼ **U**(3*a*, 3*b*), *j* ∈ **group 2**). We forced all taxa in both groups to have identical model coefficients (i.e., *β*_*i*,2_ = 0).

Finally, we consider performance when taxon-specific sequencing depth effects do not exist. We limited the sample size for each group to 500 with 50% DA taxa.

We normalized the simulated raw counts with TaxaNorm and several other methods, namely ANCOM-BC, TSS, TMM, CSS, and Wrench (Supplementary Table 3). We conducted DA testing with the post-normalized counts for each taxon using the Wilcoxon rank-sum test, adjusted p-values using the Benjamini-Hochberg method [45] for false discovery rate (FDR) control at 0.05, and compared performance in terms of power and FDR for detecting true DA taxa. We repeated each simulation scenario procedure 1000 times.

## Results

### Performance for diagnosis tests

When assessing the performance of the model diagnosis tests proposed in method section, TaxaNorm showed adequate power and good control for type I error in both prevalence and equivalence tests. When no sequencing effect exists for any taxa, the prevalence test controls the type I error at a nominal level of 5% (Table 1). Power is lacking with smaller sample sizes when the majority of taxa are not influenced by sequencing depth. However, the prevalence test showed increased power for identifying sequencing depth with larger sample sizes (Table 1). This is expected because TaxaNorm relies on a regression model, which requires a sufficient sample size for accurate parameter estimation. Specifically, with a sufficiently large sample size, power will reach 100% (Table 1). When the sequencing depth effect exists for at least one taxon but varies across taxa, the equivalence test shows high stability with power over 90% (Table 2). Under the null hypothesis, with a consistent sequencing depth effect for all taxa, the type I error rate for the equivalence test is also under 10% (Table 2). In addition, TaxaNorm continues to produce reliable parameter estimations for *β*_*i*,0_ and *β*_*i*,1_ with the EM algorithm (Supplementary Figure 6). Estimation for zero and dispersion parameters is not perfectly unbiased, but this is within expectations because these estimates are unstable for the ZINB model [42]. As shown in the simulation results for the DA analysis outlined later, downstream analysis is not affected even though the parameters are biased.

**Table 1.**
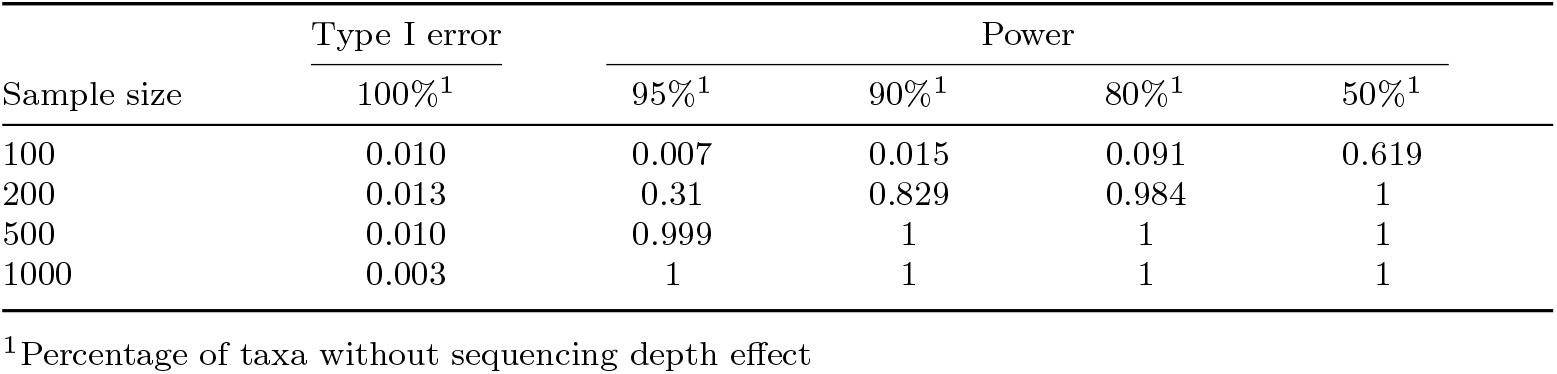
Power and type I error for prevalence test.

**Table 2.** Power and type I error for equivalence test.

| Sample size | Type I error | Power |  |  |  |
| --- | --- | --- | --- | --- | --- |
|  | 100%<sup>1</sup> | 95%<sup>1</sup> | 90%<sup>1</sup> | 80%<sup>1</sup> | 50%<sup>1</sup> |
| 100 | 0.001 | 0.988 | 0.982 | 0.983 | 0.992 |
| 200 | 0.060 | 1 | 1 | 1 | 1 |
| 500 | 0.086 | 1 | 1 | 1 | 1 |
| 1000 | 0.071 | 1 | 1 | 1 | 1 |

<sup>1</sup>Percentage of taxa with non-equal sequencing depth effect

### Performance for DA analysis

The simulation results indicate that, in various settings, TaxaNorm has better overall performance for balancing power and FDR compared to existing methods. When biological differences are small (fold change from *U* (2, 4)), only TaxaNorm, ANCOM-BC, and CSS have good control of FDR at a 5% nominal level regardless of the sample size and proportion of DA taxa (Figure 3b, 3d, and 3f). In particular, among these three methods, TaxaNorm is the most powerful when the proportion of DA taxa is smaller (Figure 3a, 3c, and 3e). TaxaNorm becomes conservative with a smaller sample size, which is within expectations, but maintains good power compared to CSS and ANCOM-BC. CSS is powerful with large sample sizes but loses a substantial amount of power when sample sizes decrease (Figure 3a, 3c). Only with 50% DA taxa does CSS have higher power than TaxaNorm. Although ANCOM-BC has the best performance for controlling FDR, its power is not as high as TaxaNorm for identifying true DA taxa. For other methods, TSS has similar power as TaxaNorm but is much more conservative with a large proportion of DA taxa. TMM has the highest power for most scenarios. Wrench and CSS have similar performance. However, all these methods have a severely inflated FDR with a range from 20% to 60% in all scenarios (Figure 3b, 3d, 3f, Supplementary Figure 2b - 4b). The FDR is uncontrolled even with larger sample sizes and fewer DA. Thus, their performance is not optimal considering the balance between power and FDR control. Interestingly, with a higher percentage of DA taxa, most methods, including TaxaNorm, lose power. However, CSS and Wrench have increased power, indicating that they may be more robust for data with greater variation. For larger biological differences (fold change from *U* (4, 10)), TaxaNorm also has compelling performance compared to existing methods (Supplementary Figure 2-4).

**Fig. 3.**
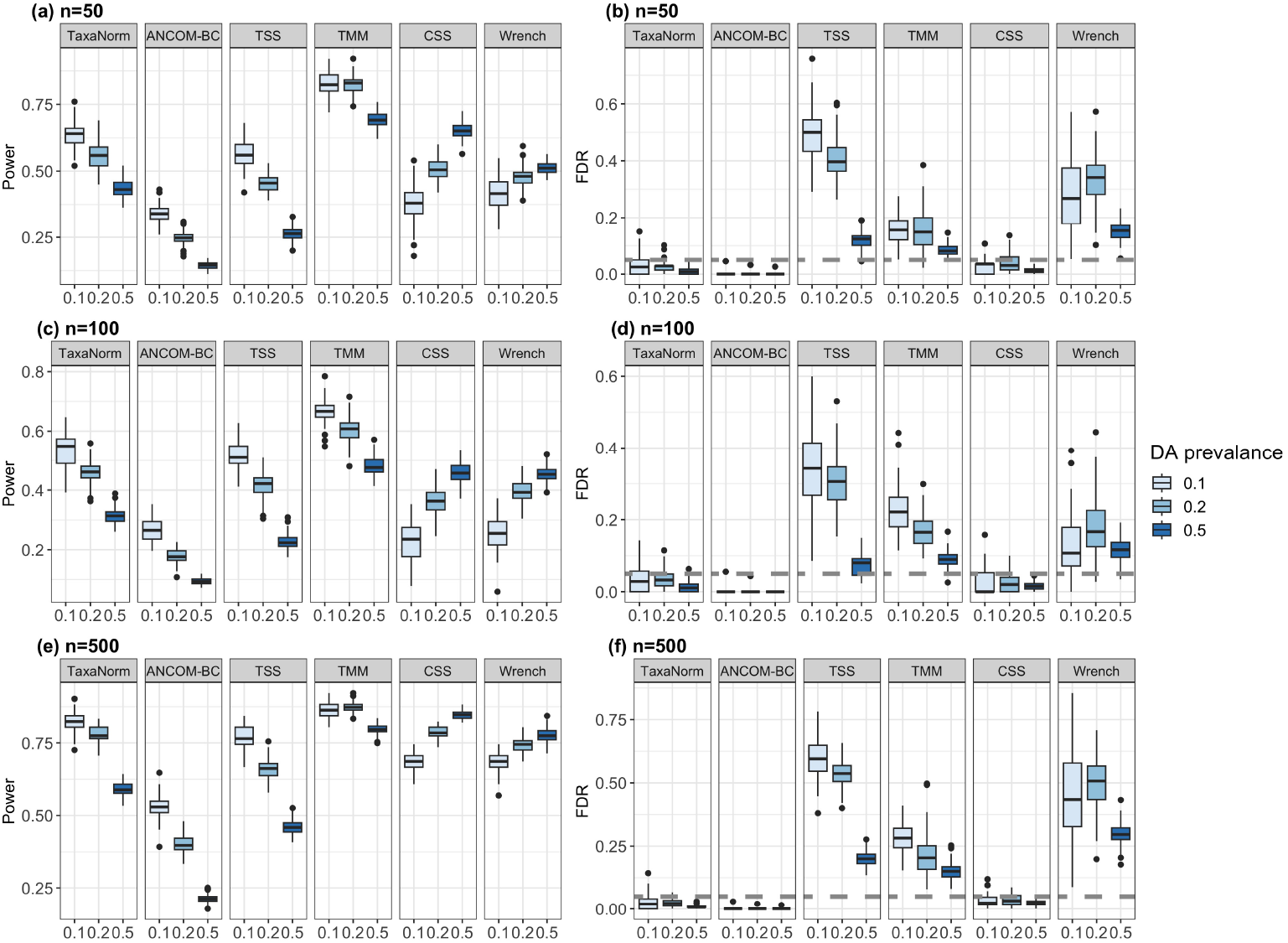
Comparison of normalization methods in terms of power and FDR in simulated datasets. The biological difference between groups is set to be small (fold-change from *U* (2, 4)). Panels (a), (c), and (e) show the power (y-axis) for identifying DA taxa with sample sizes of 50, 100, and 500 per group, and panels (b), (d), and (f) show the FDR (y-axis). In three simulation scenarios, various percentages of DA taxa (0.1, 0.2, 0.5), denoted by the colors shown in the legend, are considered. The panels are labeled by normalization method. The Benjamini-Hochberg procedure was used to adjust for the multiple testing burden with 5% as the nominal FDR level (dashed line). Number of simulations = 1000.

For datasets without DA taxa, we calculated the log fold-change values for each taxon. Ideally, in normalized data, the estimated log fold-change for post-normalized data is around 0. As shown in Figure 4, both TaxaNorm and CSS provide unbiased estimates of log fold-change with the smallest variation (Table 3). ANCOM-BC and TSS also yield unbiased estimates of log fold-changes, but with larger variations, and TMM and Wrench yield biased estimates (Figure 4, Table 3).

**Table 3.** Summary statistics for estimated fold-changes.

|  | Mean | SD<sup>1</sup> | Min | Max |
| --- | --- | --- | --- | --- |
| TaxaNorm | 0.0009 | 0.0199 | -0.0586 | 0.0662 |
| ANCOM-BC | -0.0022 | 0.0700 | -0.3169 | 0.5613 |
| TSS | 0.0009 | 0.0489 | -0.2157 | 0.4503 |
| TMM | -0.0120 | 0.0864 | -0.4006 | 0.9335 |
| CSS | -0.0006 | 0.0124 | -0.1111 | 0.0981 |
| Wrench | -0.0113 | 0.0680 | -0.3003 | 0.5372 |

<sup>1</sup>Standard deviation

**Fig. 4.**
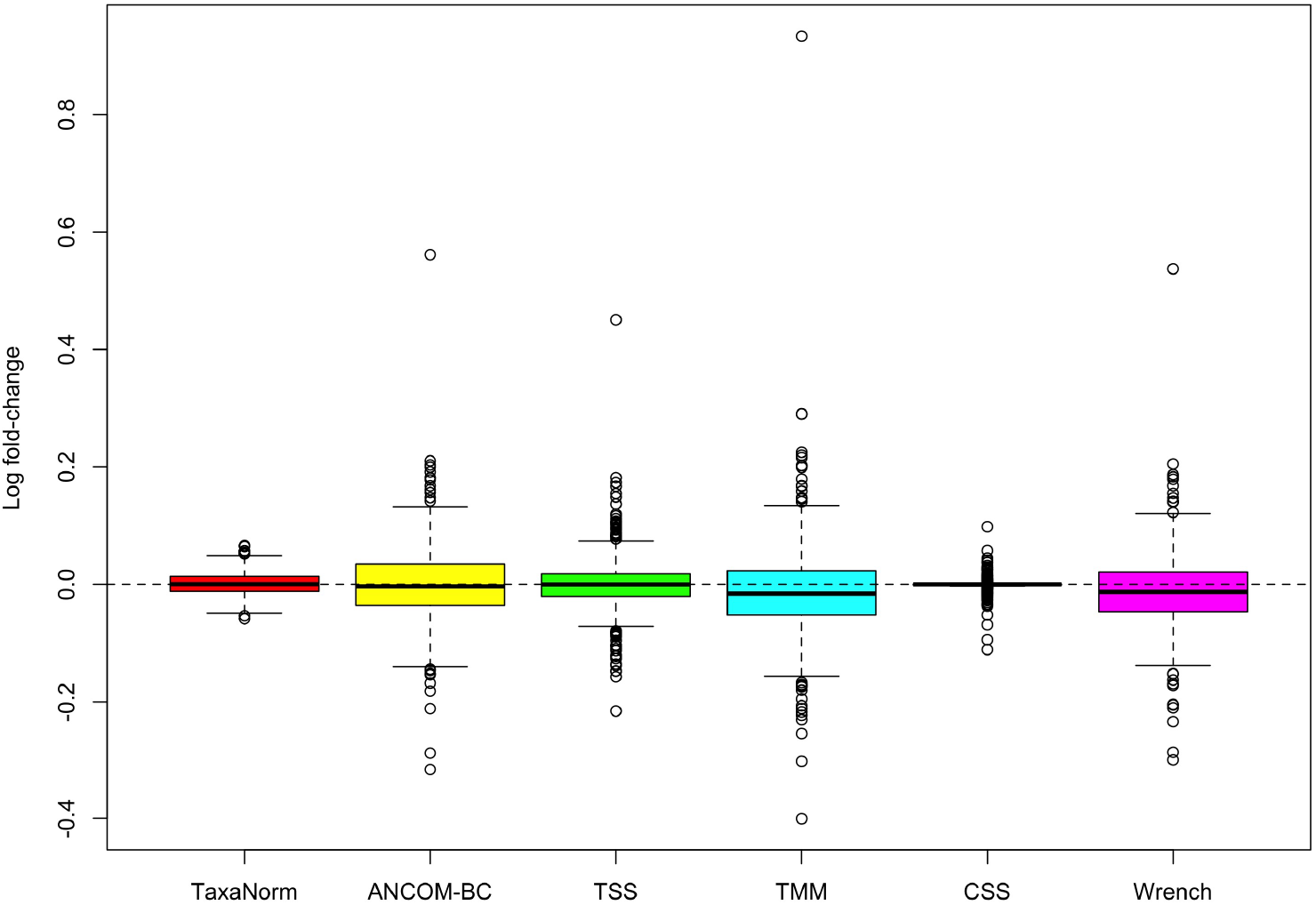
Comparison of normalization methods in terms of estimated fold-changes in simulated datasets. No true DA taxa were simulated. The boxplots show the log fold-change values for all taxa for the two simulated groups after applying each normalization method. The dashed line denotes a log fold-change value of 0. Number of simulations = 1000.

When assuming consistent sequencing efficiency for all taxa, TaxaNorm still controlled the FDR at a nominal level (5%) and maintained the ability to identify true DA taxa. Specifically, TaxaNorm was slightly less powerful than TMM, CSS, and Wrench, which is expected since these methods were developed under such an assumption (Supplementary Figure 5). Thus, TaxaNorm is robust even when the data do not satisfy our model assumption.

### Application to Human Microbiome Project data

We applied our method to normalize raw taxa counts in data from the Human Microbiome Project (HMP) [46, 47]. The samples were from a human microbiome catalog comprising samples collected from five major body sites (oral cavity, nasal cavity, skin, gastrointestinal tract, and urogenital tract) in 300 healthy individuals aged between 18 and 40 years. Subjects provided samples at up to three visits, and taxonomic profiles were generated from 16S and WGS. Reads were deposited into the Data Analysis and Coordination Center, and the taxa counts and metadata can be downloaded from https://www.hmpdacc.org/hmp/. The data collection protocol and samples are described elsewhere [46, 47]. The downloaded DA taxa data were already proceed using QIMME for 16S [38] and MetaPhlAn3 for WGS [48]. We further filtered out rare taxa with more than 10 zeros across all samples before normalization and any downstream analysis.

We first applied various normalization methods to data on the HMP gut mircobiota samples to determine their performance for eliminating the effect of sequencing depth. We examined the relationship between counts and sequencing depth before and after normalization with various methods, including TaxaNorm for all taxa. Figure 1 presents two taxa (*Dehalobacterium* and *Bacteroides*) with different abundance and sparsity levels for illustration. As expected, the raw taxa counts increased with sequencing depth, but the trends were not identical (Figure 1a). However, employing existing scaling methods with a global sample-specific size factor resulted in only partial removal of the sequencing depth effect, regardless of whether the methods were developed with RNA-seq data or microbiome data, while TaxaNorm completely removed the influence of sequencing depth on both taxa (Figure 1d). With the prevalence and equivalence tests proposed in the method section, we found sequencing depth has a non-zero effect on at least one taxon (p-value *<* 1*e*^−10^), and the effect was not identical across taxa (p-value *<* 1*e*^−10^). The results from diagnosis tests were consistent with those in Figure 2, which shows that the count-sequencing depth relationship varies by taxa. More specifically, taxa from the same phylum group had similar coefficients of sequencing depth whereas the coefficients differed across different taxonomy group. Also, although most taxa had a moderate to strong sequencing depth effect, some taxa had a less obvious effect with coefficients around zero.

We then performed non-metric multidimensional scaling (NMDS) to assess how TaxaNorm and other normalization methods influence downstream analysis to distinguish samples by phenotype. For this, we restricted our analysis to the WGS data, which consists of 1,192 taxa obtained from 749 samples from five sites on the human body, including stool (n=147), skin (n=27), vagina (n=67), oral cavity (n=415), and nasal cavity (n=93). As shown in Figure 5, TaxaNorm has good performance when visually separating samples from different body sites, particularly skin and nasal cavity samples, compared to other methods (Figure 5a). For applications of ANCOM-BC, TSS, CSS, and Wrench, the skin samples were mixed with nasal cavity samples (Figure 5b, 5c, 5e, 5f). CSS even classified skin samples into 2 sub-clusters on the NMDS2 scale (Figure 5e). TSS, TMM, CSS, and Wrench had poor performance for differentiating vagina samples and oral cavity samples (Figure 5c-f). ANCOM-BC provided a poor classification of nasal cavity and oral cavity samples (Figure 5b). All six methods had similar performance for dividing nasal cavity and vagina samples, but TaxaNorm produced a clearer separation on the NMDS2 scale. Additionally, TaxaNorm produced the largest between-group sum of squares (BSS) value and was the only method with an improved BSS value compared to the raw data (Supplementary Figure 9). TMM and CSS produced much smaller BSS values than the other methods, indicating that TMM and CSS are less optimal choices for clustering.

**Fig. 5.**
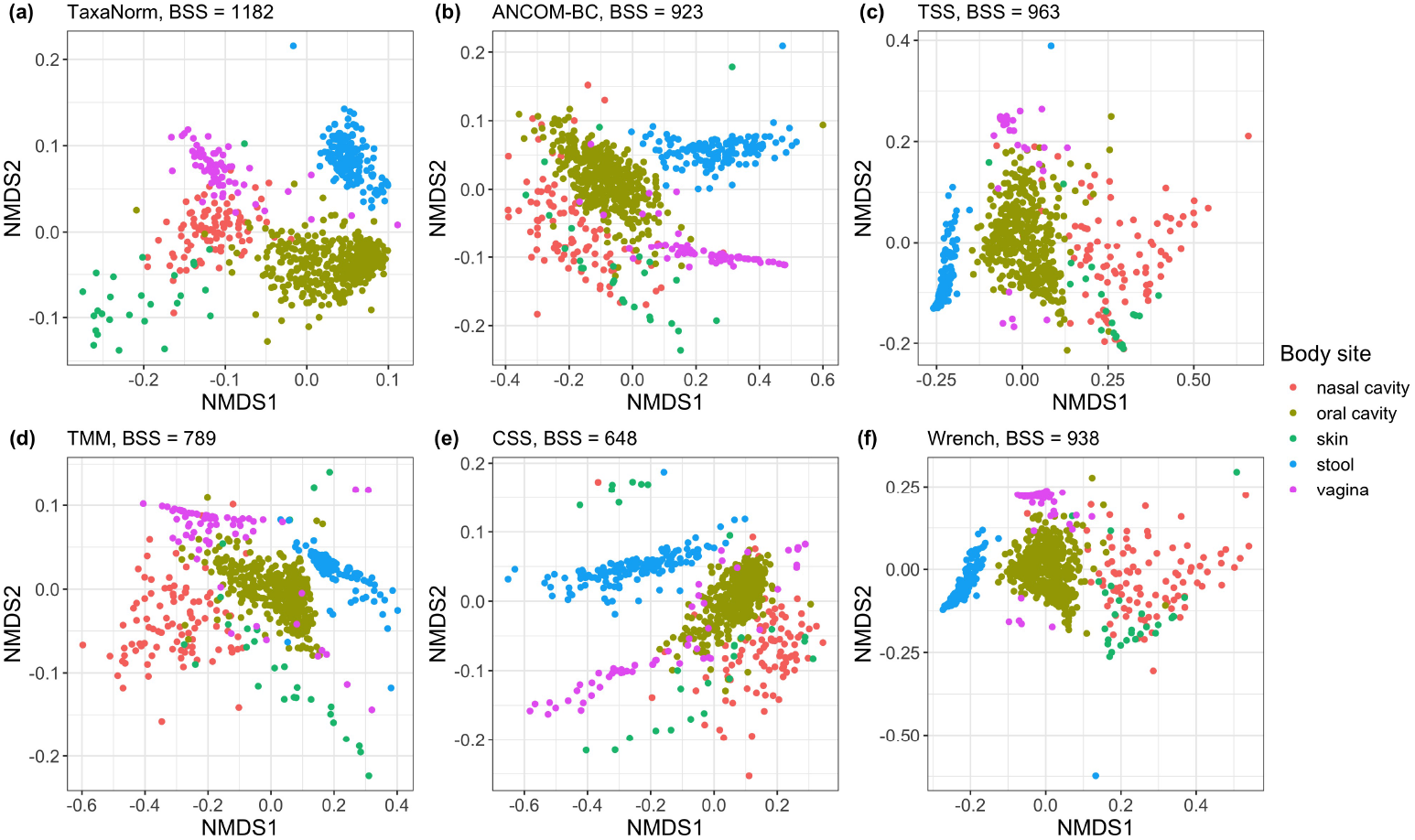
NMDS visualizations for normalized HMP data. Two NMDS coordinates were used to evaluate the performance of various normalization methods: (a) TaxaNorm, (b) ANCOM-BC, (c) TSS, (d) TMM, (e) CSS, and (f) Wrench. The sample type is denoted by the colors shown on the legend (nasal cavity, oral cavity, skin, stool, vagina). Values for between-group sum of squares (BSS) are also given.

We also report results of DA analyses for the five body sites using data normalized via TaxaNorm and other methods. We used the Benjamini-Hochberg method for multiple testing adjustment to control the FDR at 0.05. TaxaNorm identified 145 DA taxa while TSS, TMM and CSS had similar results with 146, 140 and 143 DA taxa, respectively. Interestingly, Wrench identified 150 DA taxa, which is the highest number of all the considered methods. ANCOM-BC identified 138 DA taxa, which is the lowest number of all the considered methods. This result is also consistent with simulation results that indicate ANCOM-BC is usually the least powerful of the methods. In addition, 125 DA taxa were identified by all methods. Five DA taxa were missed by TaxaNorm but identified by the other methods (Figure 6).

**Fig. 6.**
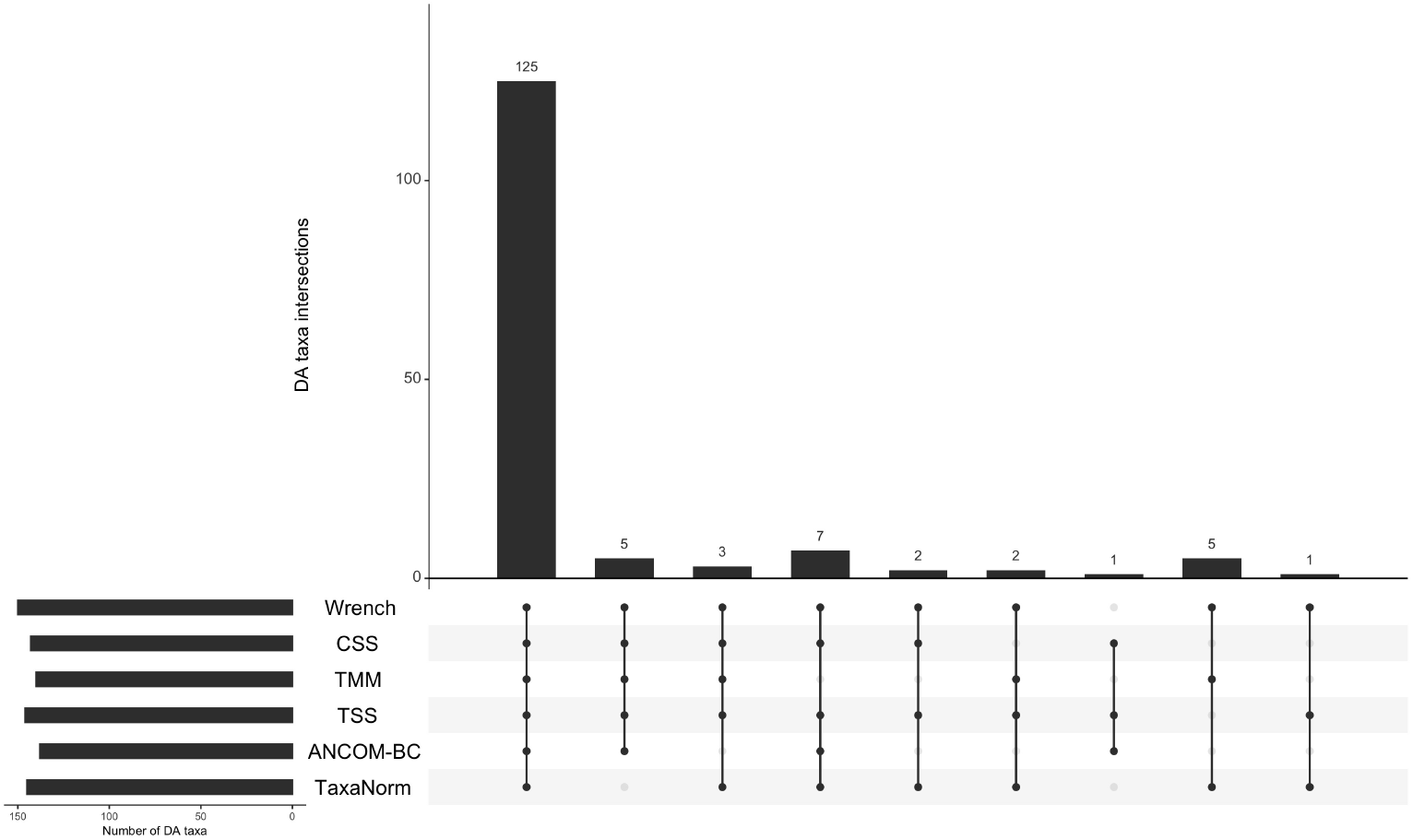
Upset plot of DA taxa determined by various normalization methods (FDR *<* 0.05). The side bar plot shows the number of DA taxa identified by each method. The main bar plot shows the intersection of DA taxa identified by multiple methods. Commonly identified DA taxa shared by various methods are aligned with vertical lines.

## Discussion

Normalizing microbiome data prior to downstream analysis is crucial because of the potential bias introduced by variations in sequencing depth across samples, which can result in undesirable and misleading conclusions regarding underlying biological mechanisms [11, 12, 16]. Normalization is conducted to remove such systematic effects so that all samples are on the same scale and independent of sequencing depth and thus the results will reflect true differences in underlying biology. However, existing normalization methods based on scaling do not sufficiently remove this effect because they violate the varying sequencing efficiency assumption and yield an elevated FDR, which results in loss of power in downstream analysis. Further, McMurdie and Holmes [12] demonstrated that other non-parametric normalization methods such as rarefaction are inadmissible and result in loss of information due to over-dispersion of the taxa count, decreasing their power.

Our proposed TaxaNorm, a novel taxa-specific normalization method for microbiome data, addresses these drawbacks and has several advantages compared to existing methods, namely taking into account the varying effects of sequencing depth across taxa. Because of its demonstrated utility in advanced sequencing experiments, we explored ZINB regression, which models taxa count with sequencing depth as a covariate and uses the residuals as normalized count values. To better differentiate the structure of excess zeros and accommodate sample heterogeneity, we finalized our model in a flexible setting by jointly modeling zero counts and sample-specific dispersion. The results of simulation studies show that TaxaNorm has good performance for identifying true DA taxa with a low FDR. In real-data applications, TaxaNorm yielded a better grouping of samples from the same biological origin (i.e., same microbial community). These results indicate that TaxaNorm offers improved accuracy and efficiency for downstream analysis and visualization compared to existing normalization methods. In addition to correcting for bias due to uneven sequencing depth, we propose two powerful tests to rigorously check the goodness-of-fit for TaxaNorm. These tests are used for model diagnosis on a per-taxon basis, and simulation results show that TaxaNorm is robust even under a non-model assumption.

Importantly, TaxaNorm is not limited to microbiome data and can be applied in any omics data produced from sequencing technologies. This functionality is particularly important in light of the recent movement to collect genomics data in epidemiologic and environmental health sciences studies. One of the limitations for TaxaNorm is that ZINB model can result in less power if the data are not truly zero-inflated. Accordingly, in practice, we incorporate a pre-possessing step to divide taxa according to their zero counts. For those with less than 5% zero counts, we instead use the varying-dispersion NB regression model (see details in the Supplementary Material). Kaul et al. [49] proposed a more sophisticated method based on differentiate structure zeros that we plan to include in our package in the future. Since TaxaNorm is built on a regression framework, its performance is also affected by sample size and outliers. For the best performance, we recommend applying our method in data with a moderate sample size and conducting winsorization for any extreme taxa counts. Considering Bayesian regression with a prior when estimating the parameters will improve the model fit. It should also be noted that the choice of bioinformatics pipelines and filtering criteria affect downstream analysis. Several benchmark papers have discussed this in depth [50–54]. However, further work should explore how TaxaNorm and other normalization methods can be customized in various situations. In future work, TaxaNorm can be also extended by including batch effects or other taxa-dependent covariates such as GC-content and genome length, which has been shown to affect taxa abundances from sequencing [9, 36, 55, 56]. Due to the flexible specification of our model setting, it is convenient and easy to include new covariates. Another potential extension is incorporating a phylogenetic tree to account for dependency between taxa. This would enable information on taxa with similar evolutionary paths to be pooled and parameters to be regularized, such that similar taxa would share the same regression coefficients in the TaxaNorm model. An intuitive method is estimating the phyla-based sequencing depth coefficients since a similar pattern was previously observed (Figure 2, Supplementary Figure 7, 8). Further, a recent paper applied the method in metatranscriptomic data and showed its feasibility [57]. This would improve computational efficiency and robustness by simplifying the model by including less parameters. Further, with the increasing popularity of longitudinal studies, the extension of TaxaNorm to mixed effects modeling on ZINB regression would be useful. Further, additional datasets could be explored to further expand and validate TaxaNorm and improve existing normalization methods.

## Conclusion

Reclaiming the true absolute feature counts (e.g., taxa abundance for microbiome data or gene expression for RNA-seq data), regardless of sequencing depth, using an advanced normalization algorithm can enable scientists to avoid deep sequencing and thus reduce the high costs associated with the technique. To address the over- or under-correcting issue identified in scaling methods, we developed a novel normalization method, TaxaNorm, to account for both sample- and taxon-specific sequencing depth effect. TaxaNorm shows improved performance with both simulated and real data and can aid in data interpretation and visualization.

## List of abbreviations

HTS: high-throughput sequencing
rRNA: ribosomal RNA
DA: differential abundance
TSS: total-sum scaling
MED: median-by-ratio
UQ: upper quartile
TMM: trimmed mean of M-values
CSS: cumulative sum scaling
ANCOM-BC: analysis of compositions of microbiomes with bias correction
ZINB: zero-inflated negative binomial
NB: negative binomial
EM: expectation-maximization
MLE: maximum likelihood estimation
PMF: probability mass function
CDF: cumulative distribution function
LRT: likelihood ratio test
FDR: false discovery rate
HMP: Human Microbiome Project
16S: 16S rRNA gene amplicon sequencing
WGS: whole-genome shotgun sequencing
NMDS: non-metric multidimensional scaling
BSS: between-group sum of squares

## Acknowledgements

The authors thank Frank Day and Deepak Mav of NIEHS for expert computational assistance. We also appreciate Shyamal D. Peddada and Thomas A. Randall for their generous suggestions on manuscript writing for NIH internal review.

## Availability of data and materials

The HMP data can be downloaded from https://www.hmpdacc.org/hmp/. The ‘TaxaNorm’ R package is freely available for download at https://github.com/wangziyue57/TaxaNorm and is available from CRAN.

## Ethics approval and consent to participate

Not applicable.

## Consent for publication

Not applicable.

## Author contributions

ZW initiated the research question, formulated the model development, conducted data analysis, and drafted the manuscript. DL created the R package. SZ and AMR provided statistical input on model specification. All authors edited and approved the final manuscript.

## Supplementary information

Supplementary methods and additional simulation results are provided.

## References

[1] Barcik, W., Boutin, R.C., Sokolowska, M., Finlay, B.B.: The role of lung and gut microbiota in the pathology of asthma. Immunity 52(2), 241–255 (2020)

[2] Cho, I., Blaser, M.J.: The human microbiome: at the interface of health and disease. Nature Reviews Genetics 13(4), 260–270 (2012)

[3] Liu, Y.-X., Qin, Y., Chen, T., Lu, M., Qian, X., Guo, X., Bai, Y.: A practical guide to amplicon and metagenomic analysis of microbiome data. Protein & Cell 12(5), 315–330 (2021)

[4] Di Bella, J.M., Bao, Y., Gloor, G.B., Burton, J.P., Reid, G.: High throughput sequencing methods and analysis for microbiome research. Journal of Microbiological Methods 95(3), 401–414 (2013)

[5] Johnson, J.S., Spakowicz, D.J., Hong, B.-Y., Petersen, L.M., Demkowicz, P., Chen, L., Leopold, S.R., Hanson, B.M., Agresta, H.O., Gerstein, M., et al.: Evaluation of 16s rRNA gene sequencing for species and strain-level microbiome analysis. Nature Communications 10(1), 1–11 (2019)

[6] Ranjan, R., Rani, A., Metwally, A., McGee, H.S., Perkins, D.L.: Analysis of the microbiome: Advantages of whole genome shotgun versus 16s amplicon sequencing. Biochemical and Biophysical Research Communications 469(4), 967–977 (2016)

[7] Quince, C., Walker, A.W., Simpson, J.T., Loman, N.J., Segata, N.: Shotgun metagenomics, from sampling to analysis. Nature Biotechnology 35(9), 833–844 (2017)

[8] Gloor, G.B., Macklaim, J.M., Pawlowsky-Glahn, V., Egozcue, J.J.: Microbiome datasets are compositional: and this is not optional. Frontiers in Microbiology 8, 2224 (2017)

[9] Sims, D., Sudbery, I., Ilott, N.E., Heger, A., Ponting, C.P.: Sequencing depth and coverage: key considerations in genomic analyses. Nature Reviews Genetics 15(2), 121–132 (2014)

[10] Quinn, T.P., Erb, I., Richardson, M.F., Crowley, T.M.: Understanding sequencing data as compositions: an outlook and review. Bioinformatics 34(16), 2870–2878 (2018)

[11] Lin, H., Peddada, S.D.: Analysis of compositions of microbiomes with bias correction. Nature Communications 11(1), 1–11 (2020)

[12] McMurdie, P.J., Holmes, S.: Waste not, want not: why rarefying microbiome data is inadmissible. PLoS Computational Biology 10(4), 1003531 (2014)

[13] Zaheer, R., Noyes, N., Ortega Polo, R., Cook, S.R., Marinier, E., Van Domselaar, G., Belk, K.E., Morley, P.S., McAllister, T.A.: Impact of sequencing depth on the characterization of the microbiome and resistome. Scientific Reports 8(1), 1–11 (2018)

[14] Pereira-Marques, J., Hout, A., Ferreira, R.M., Weber, M., Pinto-Ribeiro, I., Van Doorn, L.-J., Knetsch, C.W., Figueiredo, C.: Impact of host DNA and sequencing depth on the taxonomic resolution of whole metagenome sequencing for microbiome analysis. Frontiers in Microbiology 10, 1277 (2019)

[15] Paulson, J.N., Stine, O.C., Bravo, H.C., Pop, M.: Differential abundance analysis for microbial marker-gene surveys. Nature Methods 10(12), 1200–1202 (2013)

[16] Weiss, S., Xu, Z.Z., Peddada, S., Amir, A., Bittinger, K., Gonzalez, A., Lozupone, C., Zaneveld, J.R., Vázquez-Baeza, Y., Birmingham, A., et al.: Normalization and microbial differential abundance strategies depend upon data characteristics. Microbiome 5(1), 1–18 (2017)

[17] Hughes, J.B., Hellmann, J.J.: The application of rarefaction techniques to molecular inventories of microbial diversity. Methods in Enzymology 397, 292–308 (2005)

[18] Navas-Molina, J.A., Peralta-Sánchez, J.M., González, A., McMurdie, P.J., Vázquez-Baeza, Y., Xu, Z., Ursell, L.K., Lauber, C., Zhou, H., Song, S.J., et al.: Advancing our understanding of the human microbiome using QIIME. In: Methods in Enzymology vol. 531, pp. 371–444. Elsevier, ??? (2013)

[19] Koren, O., Knights, D., Gonzalez, A., Waldron, L., Segata, N., Knight, R., Huttenhower, C., Ley, R.E.: A guide to enterotypes across the human body: meta-analysis of microbial community structures in human microbiome datasets. PLoS Computational Biology 9(1), 1002863 (2013)

[20] Aitchison, J.: The statistical analysis of compositional data. Journal of the Royal Statistical Society: Series B (Methodological) 44(2), 139–160 (1982)

[21] Egozcue, J.J., Pawlowsky-Glahn, V., Mateu-Figueras, G., Barcelo-Vidal, C.: Isometric logratio transformations for compositional data analysis. Mathematical Geology 35(3), 279–300 (2003)

[22] Mandal, S., Van Treuren, W., White, R.A., Eggesbø, M., Knight, R., Peddada, S.D.: Analysis of composition of microbiomes: a novel method for studying microbial composition. Microbial Ecology in Health and Disease 26(1), 27663 (2015)

[23] Morton, J.T., Marotz, C., Washburne, A., Silverman, J., Zaramela, L.S., Edlund, A., Zengler, K., Knight, R.: Establishing microbial composition measurement standards with reference frames. Nature Communications 10(1), 1–11 (2019)

[24] Fernandes, A.D., Reid, J.N., Macklaim, J.M., McMurrough, T.A., Edgell, D.R., Gloor, G.B.: Unifying the analysis of high-throughput sequencing datasets: characterizing RNA-seq, 16s rRNA gene sequencing and selective growth experiments by compositional data analysis. Microbiome 2(1), 1–13 (2014)

[25] Lin, H., Peddada, S.D.: Analysis of microbial compositions: a review of normalization and differential abundance analysis. NPJ Biofilms and Microbiomes 6(1), 1–13 (2020)

[26] Costea, P.I., Zeller, G., Sunagawa, S., Bork, P.: A fair comparison. Nature Methods 11(4), 359–359 (2014)

[27] Paulson, J.N., Bravo, H.C., Pop, M.: Reply to:” a fair comparison”. Nature Methods 11(4), 359–360 (2014)

[28] Love, M.I., Huber, W., Anders, S.: Moderated estimation of fold change and dispersion for RNA-seq data with DESeq2. Genome Biology 15(12), 1–21 (2014)

[29] Bullard, J.H., Purdom, E., Hansen, K.D., Dudoit, S.: Evaluation of statistical methods for normalization and differential expression in mRNA-seq experiments. BMC Bioinformatics 11(1), 1–13 (2010)

[30] Robinson, M.D., Oshlack, A.: A scaling normalization method for differential expression analysis of RNA-seq data. Genome Biology 11(3), 1–9 (2010)

[31] Robinson, M.D., McCarthy, D.J., Smyth, G.K.: edgeR: a bioconductor package for differential expression analysis of digital gene expression data. Bioinformatics 26(1), 139–140 (2010)

[32] Kumar, M.S., Slud, E.V., Okrah, K., Hicks, S.C., Hannenhalli, S., Corrada Bravo, H.: Analysis and correction of compositional bias in sparse sequencing count data. BMC Genomics 19(1), 1–23 (2018)

[33] Gonzalez, J.M., Portillo, M.C., Belda-Ferre, P., Mira, A.: Amplification by PCR artificially reduces the proportion of the rare biosphere in microbial communities. PloS One 7(1), 29973 (2012)

[34] Wu, J.-Y., Jiang, X.-T., Jiang, Y.-X., Lu, S.-Y., Zou, F., Zhou, H.-W.: Effects of polymerase, template dilution and cycle number on PCR based 16s rRNA diversity analysis using the deep sequencing method. BMC Microbiology 10(1), 1–7 (2010)

[35] Wintzingerode, F., Göbel, U.B., Stackebrandt, E.: Determination of microbial diversity in environmental samples: pitfalls of PCR-based rRNA analysis. FEMS Microbiology Reviews 21(3), 213–229 (1997)

[36] McLaren, M.R., Willis, A.D., Callahan, B.J.: Consistent and correctable bias in metagenomic sequencing experiments. eLife 8, 46923 (2019)

[37] Lin, H., Eggesbø, M., Peddada, S.D.: Linear and nonlinear correlation estimators unveil undescribed taxa interactions in microbiome data. Nature Communications 13(1), 1–16 (2022)

[38] Schiffer, L., Azhar, R., Shepherd, L., Ramos, M., Geistlinger, L., Huttenhower, C., Dowd, J.B., Segata, N., Waldron, L.: Hmp16sdata: efficient access to the human microbiome project through bioconductor. American journal of epidemiology 188(6), 1023–1026 (2019)

[39] Pop, M., Walker, A.W., Paulson, J., Lindsay, B., Antonio, M., Hossain, M.A., Oundo, J., Tamboura, B., Mai, V., Astrovskaya, I., et al.: Diarrhea in young children from low-income countries leads to large-scale alterations in intestinal microbiota composition. Genome biology 15, 1–12 (2014)

[40] Chen, J., Chia, N., Kalari, K.R., Yao, J.Z., Novotna, M., Paz Soldan, M.M., Luckey, D.H., Marietta, E.V., Jeraldo, P.R., Chen, X., et al.: Multiple sclerosis patients have a distinct gut microbiota compared to healthy controls. Scientific Reports 6(1), 28484 (2016)

[41] Scher, J.U., Sczesnak, A., Longman, R.S., Segata, N., Ubeda, C., Bielski, C., Rostron, T., Cerundolo, V., Pamer, E.G., Abramson, S.B., et al.: Expansion of intestinal prevotella copri correlates with enhanced susceptibility to arthritis. eLife 2, 01202 (2013)

[42] Jiang, R., Zhan, X., Wang, T.: A flexible zero-inflated poisson-gamma model with application to microbiome sequence count data. Journal of the American Statistical Association 118(542), 792–804 (2023)

[43] Silverman, J.D., Roche, K., Mukherjee, S., David, L.A.: Naught all zeros in sequence count data are the same. Computational and Structural Biotechnology Journal 18, 2789–2798 (2020)

[44] Dunn, P.K., Smyth, G.K.: Randomized quantile residuals. Journal of Computational and Graphical Statistics 5(3), 236–244 (1996)

[45] Benjamini, Y., Hochberg, Y.: Controlling the false discovery rate: a practical and powerful approach to multiple testing. Journal of the Royal Statistical Society: Series B (Methodological) 57(1), 289–300 (1995)

[46] The Human Microbiome Project Consortium: A framework for human microbiome research. Nature 486, 215–221 (2012) 10.1038/nature11209

[47] The Human Microbiome Project Consortium: Structure, function and diversity of the healthy human microbiome. Nature 486(7402), 207–214 (2012)

[48] Pasolli, E., Schiffer, L., Manghi, P., Renson, A., Obenchain, V., Truong, D.T., Beghini, F., Malik, F., Ramos, M., Dowd, J.B., et al.: Accessible, curated metagenomic data through experimenthub. Nature methods 14(11), 1023–1024 (2017)

[49] Kaul, A., Mandal, S., Davidov, O., Peddada, S.D.: Analysis of microbiome data in the presence of excess zeros. Frontiers in Microbiology 8, 2114 (2017)

[50] Cao, Q., Sun, X., Rajesh, K., Chalasani, N., Gelow, K., Katz, B., Shah, V.H., Sanyal, A.J., Smirnova, E.: Effects of rare microbiome taxa filtering on statistical analysis. Frontiers in microbiology 11, 607325 (2021)

[51] Lindgreen, S., Adair, K.L., Gardner, P.P.: An evaluation of the accuracy and speed of metagenome analysis tools. Scientific reports 6(1), 19233 (2016)

[52] Simon, H.Y., Siddle, K.J., Park, D.J., Sabeti, P.C.: Benchmarking metagenomics tools for taxonomic classification. Cell 178(4), 779–794 (2019)

[53] Vollmers, J., Wiegand, S., Kaster, A.-K.: Comparing and evaluating metagenome 18 assembly tools from a microbiologist’s perspective-not only size matters! PloS one 12(1), 0169662 (2017)

[54] McIntyre, A.B., Ounit, R., Afshinnekoo, E., Prill, R.J., Hénaff, E., Alexander, N., Minot, S.S., Danko, D., Foox, J., Ahsanuddin, S., et al.: Comprehensive bench-marking and ensemble approaches for metagenomic classifiers. Genome biology 18, 1–19 (2017)

[55] Browne, P.D., Nielsen, T.K., Kot, W., Aggerholm, A., Gilbert, M.T.P., Puetz, L., Rasmussen, M., Zervas, A., Hansen, L.H.: GC bias affects genomic and metagenomic reconstructions, underrepresenting gc-poor organisms. GigaScience 9(2), 008 (2020)

[56] Ross, M.G., Russ, C., Costello, M., Hollinger, A., Lennon, N.J., Hegarty, R., Nusbaum, C., Jaffe, D.B.: Characterizing and measuring bias in sequence data. Genome Biology 14(5), 1–20 (2013)

[57] Klingenberg, H., Meinicke, P.: How to normalize metatranscriptomic count data for differential expression analysis. PeerJ 5, 3859 (2017)

