## Supplementary file for "TaxaNorm: a novel taxa-specific normalization approach for microbiome data"

### 1 Details of TaxaNorm framework

#### 1.1 ZINB distribution

Let $f_{nb}(y;\mu,\theta)$ denote the PMF of the NB distribution. The PMF of ZINB distribution is given by

$$\begin{aligned} f_{zinb}(y_{ij};\mu_{ij},\theta_{ij},\pi_{ij}) &= \pi_{ij}\cdot \mathbf{I}\left(y_{ij}=0\right)+(1-\pi_{ij})\cdot f_{nb}(y_{ij};\mu_{ij},\theta_{ij}) \\ &= \pi_{ij}\cdot \mathbf{I}\left(y_{ij}=0\right)+(1-\pi_{ij})\cdot \frac{\Gamma(y_{ij}+\theta_{ij})}{\Gamma(y_{ij}+1)\Gamma(\theta_{ij})}\left(\frac{\mu_{ij}}{\theta_{ij}+\mu_{ij}}\right)^{y_{ij}}\left(\frac{\theta_{ij}}{\theta_{ij}+\mu_{ij}}\right)^{\theta_{ij}}, \end{aligned} \tag{1}$$

#### 1.2 MLE for NB regression

When zero-inflation is not assessed from the data, a varying-dispersion NB regression can be used instead, by forcing $\pi_{ij}=0$. Then, let $\Lambda=(\beta_{i,0},\beta_{i,1},\kappa_{i,0},\kappa_{i,1})$ represent the full set of unknown parameters. The log-likelihood can be written as follows:

$$l(\Lambda;Y_i)=\sum_{j=1}^{n}\log\left\{\frac{\Gamma(y_{ij}+\theta_{ij})}{\Gamma(y_{ij}+1)\Gamma(\theta_{ij})}\left(\frac{\mu_{ij}}{\theta_{ij}+\mu_{ij}}\right)^{y_{ij}}\left(\frac{\theta_{ij}}{\theta_{ij}+\mu_{ij}}\right)^{\theta_{ij}}\right\}, \tag{2}$$

The MLE $\hat{\Lambda}=(\hat{\beta}_{i,0},\hat{\beta}_{i,1},\hat{\kappa}_{i,0},\hat{\kappa}_{i,1})$ can be obtained by directly maximizing the above Equation.

#### 1.3 TaxaNorm workflow

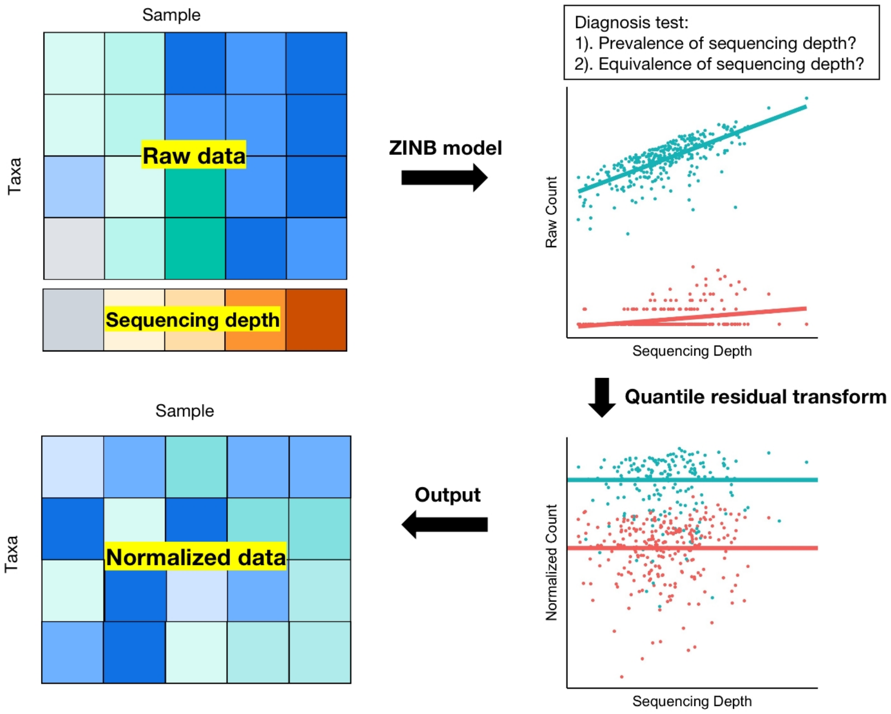


Figure 1: Overview of the TaxaNorm workflow. The input data are the raw taxa count matrix and sequencing depth vector. A varying dispersion ZINB model was used to estimate parameters for each taxon. Two diagnosis tests (prevalence and equivalence) were applied to assess taxa characteristics. Quantile residuals were used as the normalized count.

### 2 Supplemental tables

#### 2.1 Details of simulation settings

| | **Prevalence test** **% of taxa with** $\beta_{i,1} = \kappa_{i,1} = 0$ | **Equivalence test** **% of taxa with** $\beta_1 = 1$ | **Sample size** |
|---|---|---|---|
| **Scenario** | 50%<br>80%<br>90%<br>95%<br>100% | 50%<br>80%<br>90%<br>95%<br>100% | 100<br>200<br>500<br>1000 |

Table 1: Simulation scenarios for model diagnosis tests.

#### 2.2 Summary of scaling methods for comparison

The observed taxa count is divided by a size factor so that sequencing depth does not confound the reconstructed taxa count. A constant size factor is generally used for all taxa within a sample but varies by sample. Let $Y_{ij}$ be the observed count for taxon $i (i = 1, ..., p)$ in sample $j (j = 1, ..., n)$ and $s_j$ denote the sample-specific size factors. The normalized taxa count is calculated as $Y_{ij}^{\text{norm}} = \frac{Y_{ij}}{s_j}$. Table 3 lists the scaling normalization methods used for comparison in this paper.

| | % of DA taxa | Fold change | Sample size per group |
|---|---|---|---|
| **Scenario** | 10% | U(2,4) | 50 vs. 50 |
| | 20% | U(4,10) | 100 vs. 100 |
| | 50% | | 500 vs. 500 |

Table 2: Simulation scenarios for DA analysis.

| Method | Estimate of $s_j$ |
|---|---|
| ANCOM-BC | $\frac{1}{p}\sum_i \left(\log X_{ij} - Z_j^\top \hat{\beta}_{\mathrm{i}}\right)$ |
| TSS | $\sum_j Y_{ij}$ |
| TMM | $\log_2(s_j) = \frac{\sum_{i\in G_*} w_{ij} M_{ij}}{\sum_i w_{ij}}$ |
| | $M_{ij} = \log_2\left(\frac{X_{ij}}{D_j}\right) - \log_2\left(\frac{X_{ir}}{D_r}\right)$ |
| | $w_{ij} = \frac{D_j - X_{ij}}{D_j X_{ij}} + \frac{D_r - X_{ir}}{D_r X_{ir}}$ |
| | $r$ is the reference sample |
| | $G^*$ represents a set of taxa that were not considered as extreme data |
| CSS | $\sum_{i:X_{ij}\le q_j^l} X_{ij}$ |
| Wrench | $\frac{1}{p}\sum_i W_{ij}\frac{q_{ij}}{q_{+i}}$ |

Table 3: Summary of scaling normalization methods

### 3 Additional simulation results

#### 3.1 Large fold change between two simulated groups

##### 3.1.1 Sample size = 50

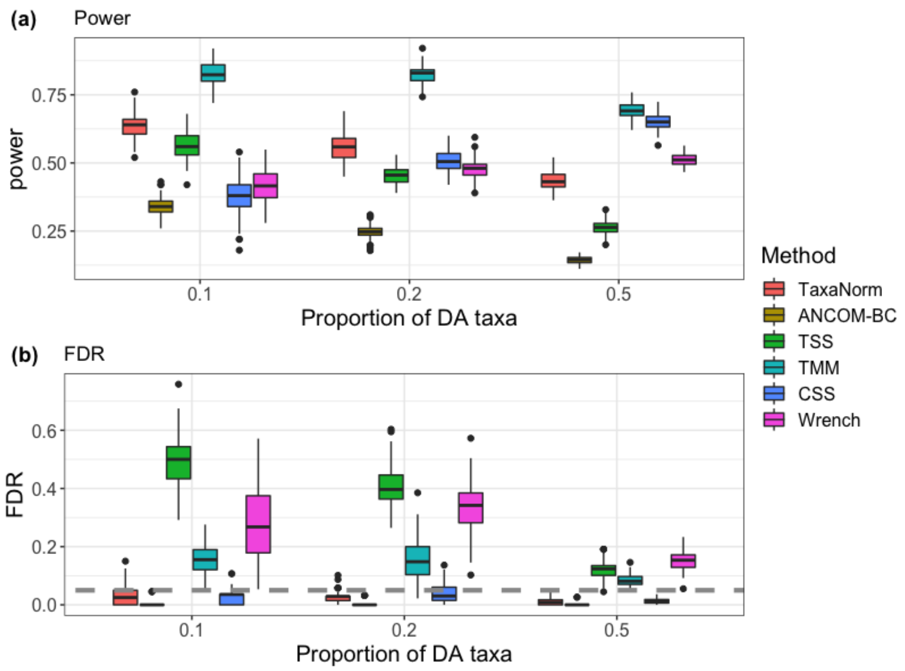


Figure 2: Comparison of different methods in terms of **(a)** power and **(b)** FDR in identifying DA taxa using simulation datasets. The biological difference between groups is set to large (fold-change from $U(4, 10)$). Each panel shows a simulation scenario with various percentages of DA taxa on the x-axis. The corresponding methods are denoted by the colors shown at the side. The BH procedure is used to adjust for multiple testing with 5% as the nominal FDR level (dashed line).

##### 3.1.2 Sample size = 100

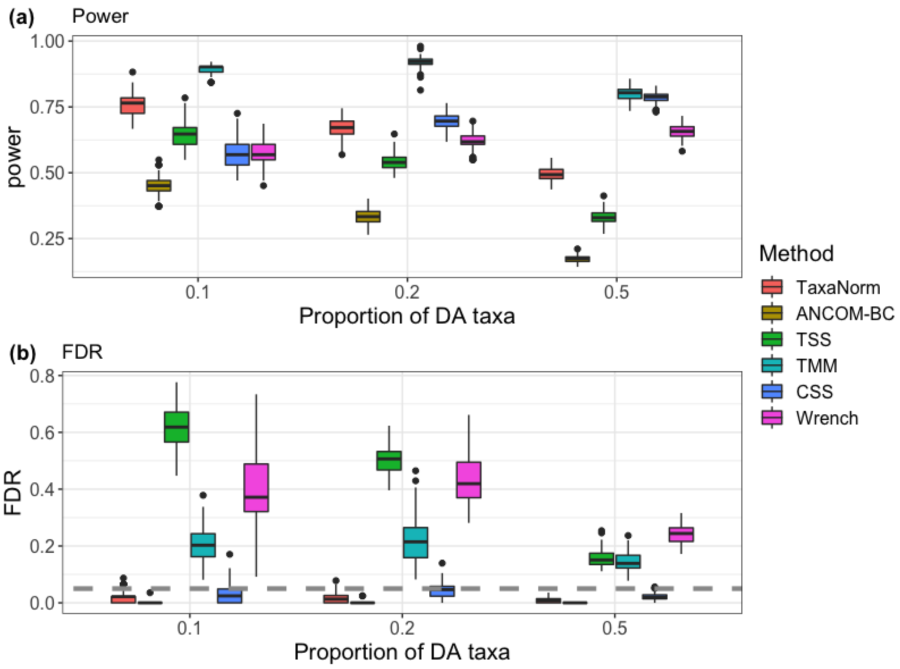


Figure 3: Comparison of different methods in terms of **(a)** power and **(b)** FDR in identifying DA taxa using simulation datasets. The biological difference between groups is set to large (fold-change from $U(4, 10)$). Each panel shows a simulation scenario with various percentages of DA taxa on the x-axis. The corresponding methods are denoted by the colors shown at the side. The BH procedure is used to adjust for multiple testing with 5% as the nominal FDR level (dashed line).

###### 3.1.3 Sample size = 500

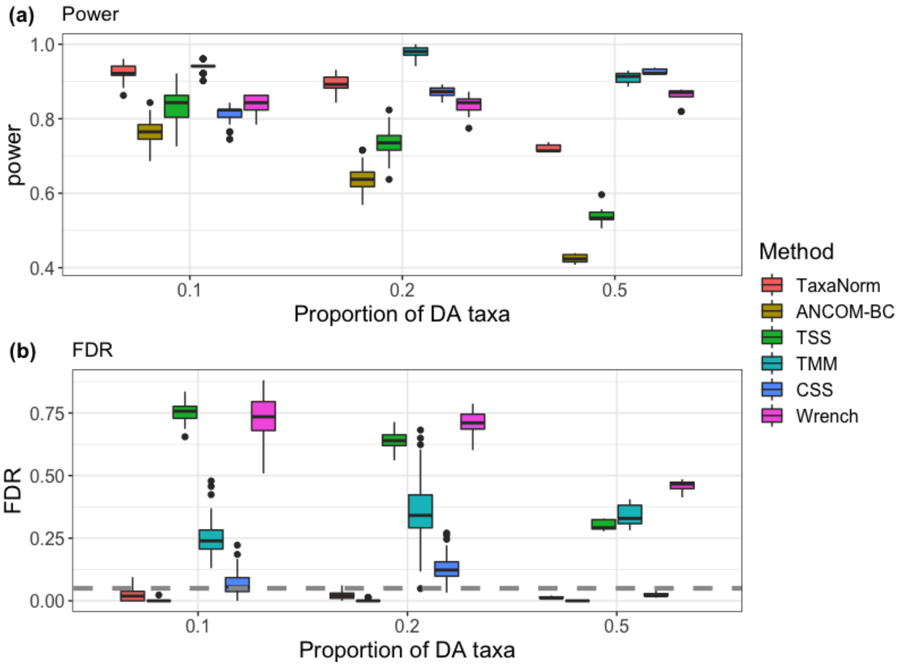


Figure 4: Comparison of different methods in terms of **(a)** power and **(b)** FDR in identifying DA taxa using simulation datasets. The biological difference between groups is set to large (fold-change from $U(4, 10)$). Each panel shows a simulation scenario with various percentages of DA taxa on the x-axis. The corresponding methods are denoted by the colors shown at the side. The BH procedure is used to adjust for multiple testing with 5% as the nominal FDR level (dashed line).

#### 3.2 Taxon-specific effect does not exist

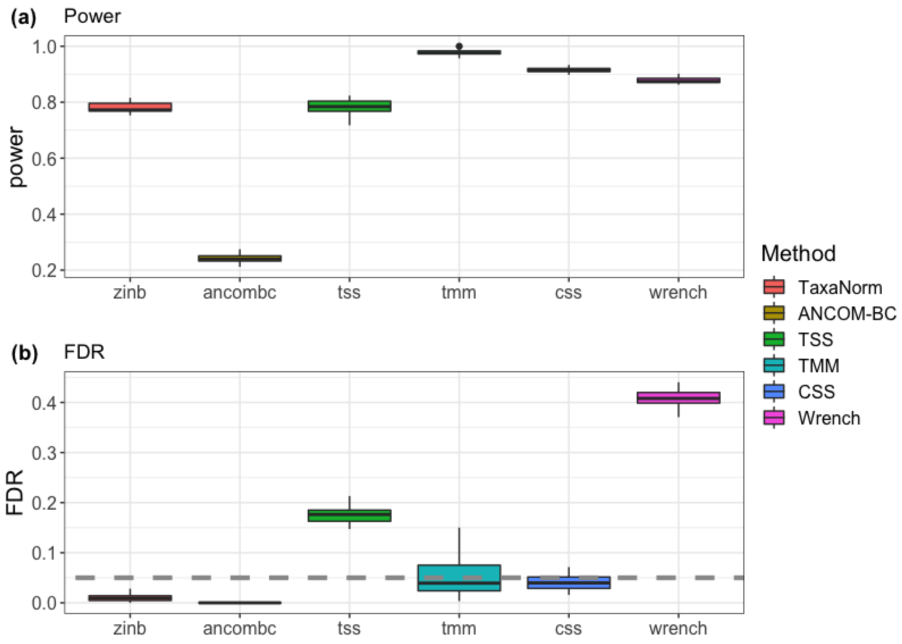


Figure 5: Comparison of different methods in terms of **(a)** power and **(b)** FDR in identifying DA taxa using simulation datasets. Sequencing effect effect are equal across all taxa. The sample size for each group is 500 with 50% DA taxa. The corresponding methods are denoted by the colors shown at the side. The BH procedure is used to adjust for multiple testing with 5% as the nominal FDR level (dashed line).

### 4 QQ plot for parameter estimation

Below are QQ-plots comparing the estimated parameters with the true values in a simulated dataset.

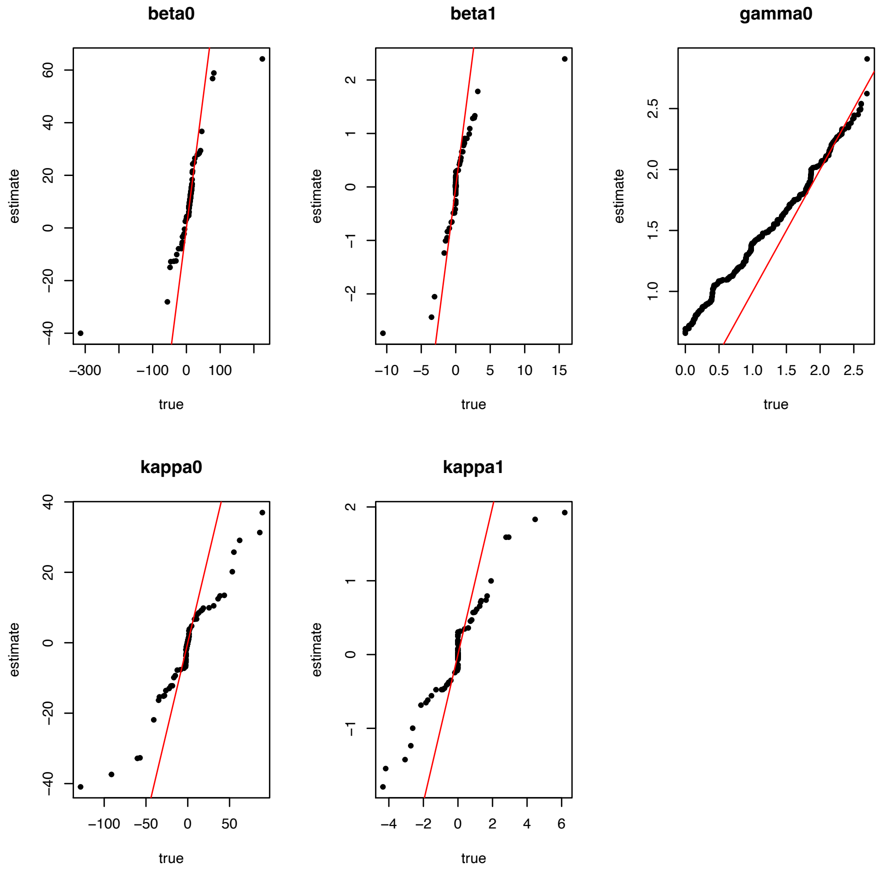


Figure 6: QQ plot for for real vs. estimated parameters in a simulation dataset with 90% of taxa without the sequencing depth effect and a sample size of 1000.

### 5 Additional real data results

#### 5.1 Count-sequencing depth relationship

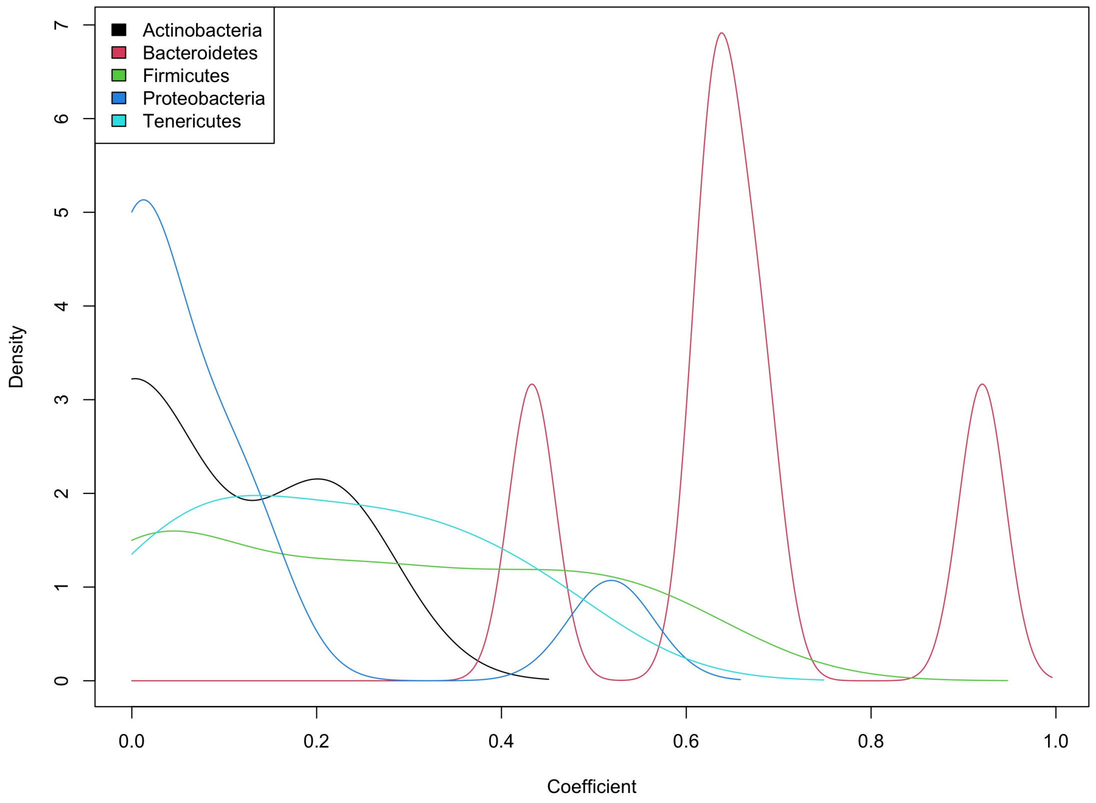


Figure 7: Sequencing efficiency of each taxon in HMP 16S gut microbiota data. The relationship between raw taxa-specific counts and sequencing depth was estimated using ZINB regression. Densities of corresponding coefficients are colored by phylum rank for all taxa.

#### 5.2 NMDS for raw HMP WGS data

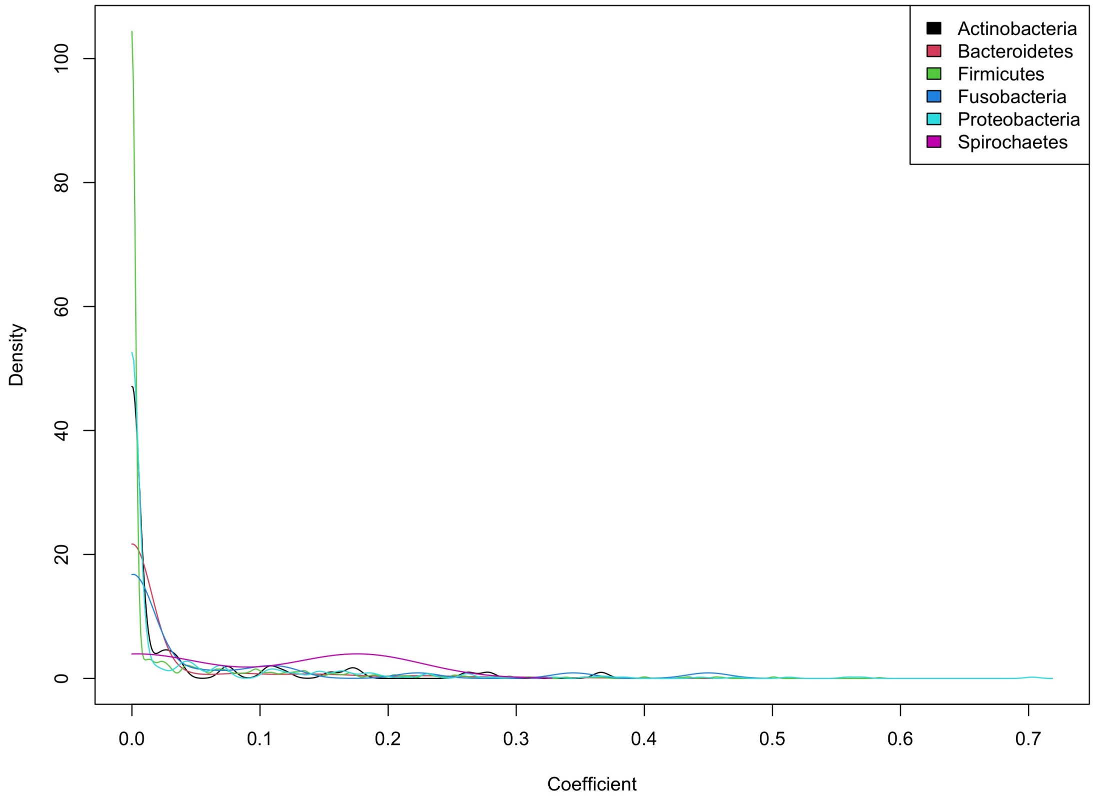


Figure 8: Sequencing efficiency of each taxon of fecal microbiota data from a diarrhea cohort. The relationship between raw taxa-specific counts and sequencing depth was estimated using ZINB regression. Densities of corresponding coefficients are colored by phylum rank for all taxa.

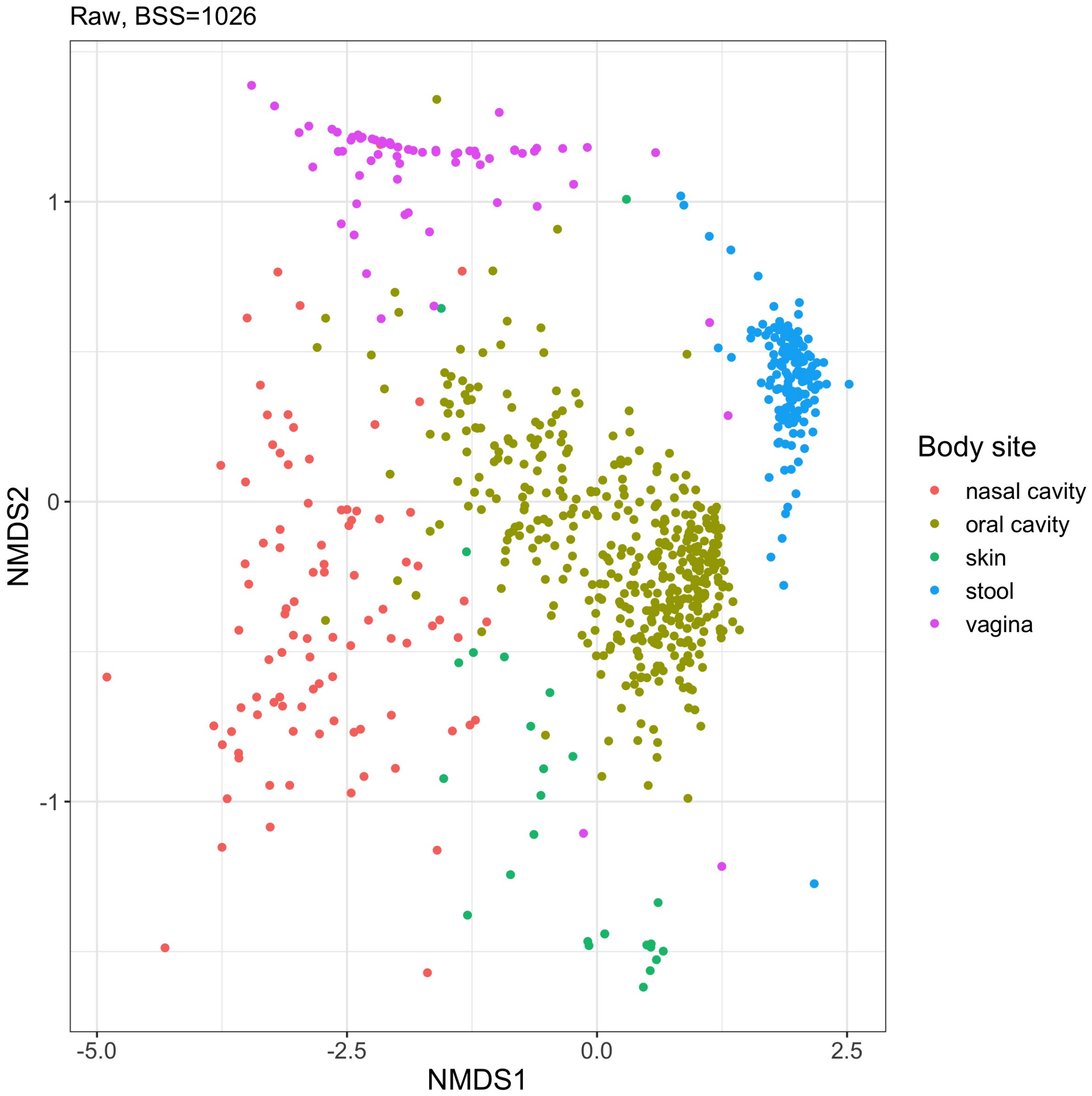


Figure 9: NMDS visualizations for raw HMP data. Two NMDS coordinates were used to present the data with BSS=1026.
